# Microfluidic study of effects of flow velocity and nutrient concentration on biofilm accumulation and adhesive strength in microchannels

**DOI:** 10.1101/375451

**Authors:** Na Liu, Tormod Skauge, David Landa-Marbán, Beate Hovland, Bente Thorbjørnsen, Florin Adrain Radu, Bartek Florczyk Vik, Thomas Baumann, Gunhild Bødtker

## Abstract

**Abstract:** Biofilm accumulation in the porous media can cause plugging and change many physical properties of porous media. Up to now, applications of desired biofilm growth and its
subsequent bioplugging have been attempted for various practices. A deeper understanding of the relative influences of hydrodynamic conditions including flow velocity and nutrient concentration, on biofilm growth and detachment is necessary to plan and analyze bioplugging experiments and field trials. The experimental results by means of microscopic imaging over a T-shape microchannel show that flow velocity and nutrient concentrations can have significant impacts on biofilm accumulation and adhesive strength in both flowing and stagnant microchannels. Increase in fluid velocity could facilitate biofilm growth, but that above a velocity threshold, biofilm detachment and inhibition of biofilm formation due to high shear stress were observed. High nutrient concentration prompts the biofilm growth, but was accompanied by a relatively weak adhesive strength. This research provides an overview of biofilm development in a hydrodynamic environment for better predicting and modelling the bioplugging associated with porous system in petroleum industry, hydrogeology, and water purification.

**IMPORTANCE:** In the recent decade, the use of bacteria has become more and more important in many applications. Bioplugging caused by bacteria growth in porous media has been explored as a viable technique for some applications, such as bioremediation, water purification and microbial enhanced oil recovery (MEOR). In order to control biofilms/biomasses selectively/directionally plugging in desirable places, the role of hydrodynamic conditions on biofilm growth and detachment is essential to investigate. Herein, a T-shape microchannel was prepared to study effects of flow velocity and nutrient concentration on biofilm accumulation and adhesive strength at pore scale. Our results suggest that flow velocity and nutrient concentration could control biofilm accumulation in microchannels. The finding helps explain and predict the engineering bioplugging in porous media, especially for the selective plugging strategy of a MEOR field trial.

## INTRODUCTION

Biofilm accumulation in the pore space can cause pore plugging, leading to significant changes in physical properties of porous media, such as the reduction of porosity and permeability (1-5). The plugging effect might have negative impacts in many industrial and medical applications because the clogging pores need extra cost to clean and prevention. However, engineering bioplugging has been explored as a viable technique for various practices, such as in situ bioremediation (6), soil injection (7), waste treatment (8, 9), ground water recharge (10) and microbial enhanced oil recovery (11-15). For example, in MEOR trails biofilm accumulation leads to selective plugging of high permeability zones, subsequently forcing the diversion of injected fluids towards lower permeable zones to improve the oil recovery (15, 16). Suthar et al. confirmed the obtained oil recovery because of bacterial growth and biofilm formation in the sand pack (17). Karambeigi et al. used two different heterogeneous micromodels to observe potential of bioplugging of high permeable layers of porous media for improving the efficiency of water flooding (2). Klueglein et al. investigated the influences of nutrients concentrations on cell growth and bioplugging potential during a MEOR trial (18). Even tremendous efforts have been made to improve the understanding of bioplugging, few works concern biofilm studies of biofilm growth and detachment mechanisms accompanying the bioplugging process.

Bioplugging in porous media results from the accumulation of bacterial cells, production of extracellular polymeric substances (EPSs) in the pore space. Due to physicochemical properties of EPSs, biofilms can behave as viscous liquids to resist the flow-induced shear stress, and substantially plug the pore (19-22). Engineering bioplugging is a process used to control biofilms selectively and substantially plugging in desired places (6, 23, 24). Therefore, knowledge on mechanisms of biofilm development and its adhesive strength with solids surface is vitally important to plan and predict the engineering bioplugging process. It was found that biofilm growth and detachment could be significantly influenced by varying hydrodynamic conditions on the surrounding environment (19, 20, 25). Biofilm growth and detachment rates could both increase with fluid velocity, as the increased mass transfer facilitating nutrients supply for bacteria growth, while the increased shear force in turn causing detachment (19, 21, 26, 26).

There is a consensus that biofilm growth rate increases with nutrients concentration, while nutrient starvation results in biofilm detachment (28-30). Nonetheless, knowledge on bioplugging must be depicted by examining a correlation between biofilm accumulations and its adhesive strength and hydrodynamic conditions like flow velocity and nutrient concentration, to improve understanding and hence ability to control bioplugging in fluid flooded porous systems.

Traditionally quiescent experiments for biofilm research were normally carried on homogeneous physical conditions, which lack environmental complexities for accurately determining the dynamic changes occurring during biofilm development (31). The advent of new technologies, specially microfluidics, have attracted a rapidly growing interest to emulate biological phenomena by addressing unprecedented control over the flow conditions, providing identical and reproducible culture conditions, as well as real-time observation (26, 30, 32, 32). Indeed, microfluidics has been used for observing biofilm formation under various fundamental and applied researches, e.g. wastewater treatment (34) and medical fields (20, 35). Herein, we used a T-shape microfluidic device equipped with a microscope to study the biofilm accumulation and adhesive strength as responds to various flow velocities and nutrient concentrations in the microchannel.

## RESULTS

### Effects of flow velocity on biofilm accumulation and adhesive strength

Biofilms development in microchannels were measured by varying injecting flowrates of 10mM pyruvate (1.0 N) from 0.2 μl/min to 0.5 μl/min. After 6 days, the shear rate was steadily increased to 500.00 s^-1^ to test the adhesive strength of biofilm attached on the solid surface. The corresponding flow velocity, Peclet number, Reynolds number and shear rate at each flowrate in Channel 1 are listed in Table 1.The accumulation of biofilms at different velocities was observed and registered as function of time by use of microscope.

**TABLE 1.**
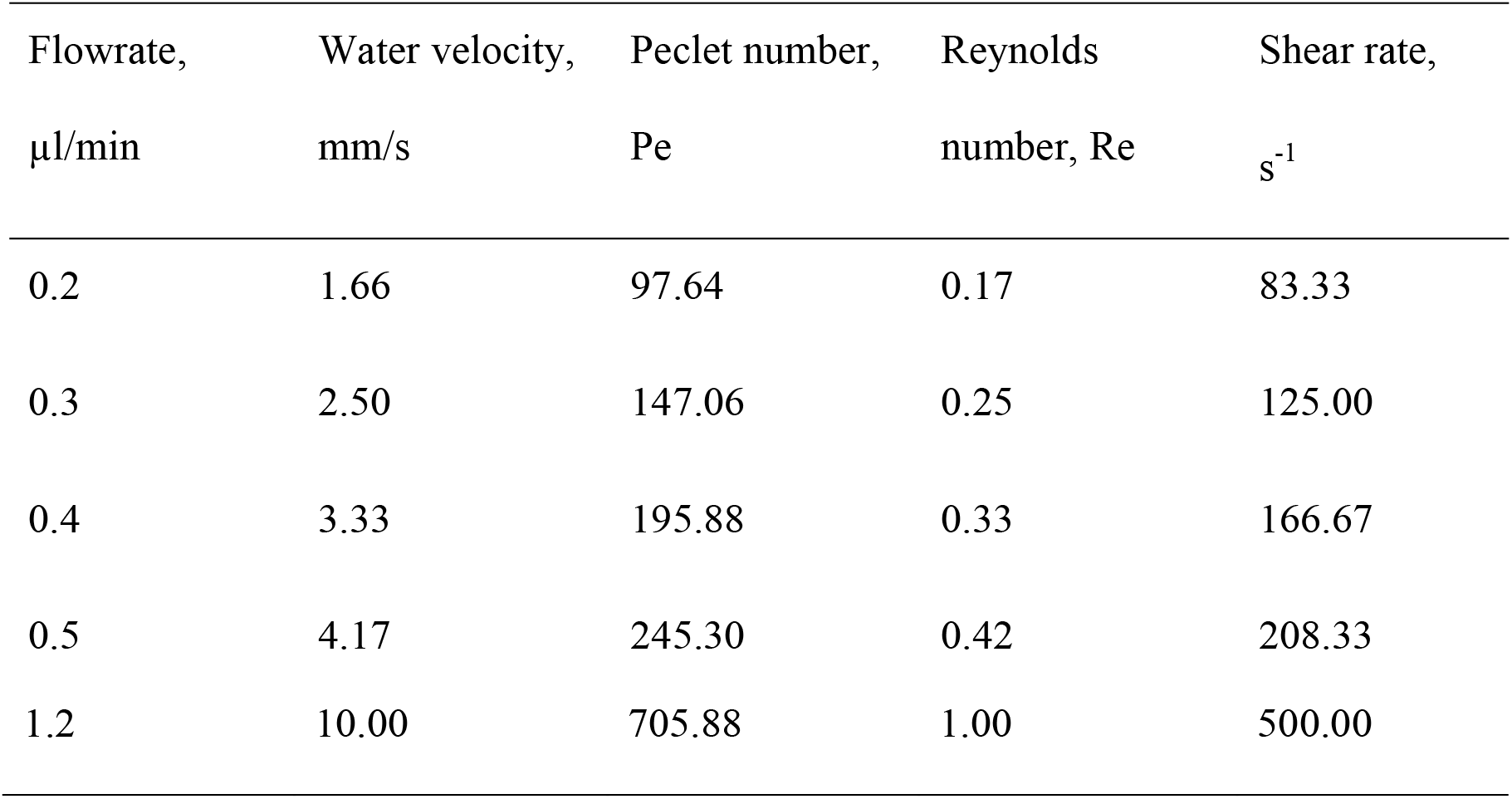
Table of basic flow parameters at various flowrates in this study.

| Flowrate,<br>$\mu\text{l}/\text{min}$ | Water velocity,<br>$\text{mm}/\text{s}$ | Peclet number,<br>Pe | Reynolds<br>number, Re | Shear rate,<br>$\text{s}^{-1}$ |
| --- | --- | --- | --- | --- |
| 0.2 | 1.66 | 97.64 | 0.17 | 83.33 |
| 0.3 | 2.50 | 147.06 | 0.25 | 125.00 |
| 0.4 | 3.33 | 195.88 | 0.33 | 166.67 |
| 0.5 | 4.17 | 245.30 | 0.42 | 208.33 |
| 1.2 | 10.00 | 705.88 | 1.00 | 500.00 |

### Biofilm morphologies

Images of biofilms development in two microchannels at various flow velocities are shown in Fig. 2. After inoculation, the initial attached biofilms at low velocities (1.66 and 2.50 mm/s) became irreversible and developed towards different structures along the nutrients flow. Biofilms at 1.66 mm/s tends to be approximately circular shape and has a larger coverage area, while biofilms at 2.50 mm/s show appearance of thin plate structures. There is no clear biofilm formation in Channel 1 at high velocities. On the contrary, biofilms formed at Channel 2 at 3.33 mm/s led to a larger clusters compared with low rates. There was no biofilm growth in either channel at the highest flow velocity of 4.17 mm/s.

After 6 days of biofilm culturing, the shear rate steadily increased to 500.00 s^-1^ to test the adhesive strength between biofilms and solid surfaces. As shown in the right column images of Fig. 2, biofilms in Channel 1 at 1.66 mm/s became elongated in the flowing direction to form filamentous “streamers” when the shear force acting on biofilms increasing with shear rate; while there was no clear shape difference on biofilms growth at higher velocities in Channel 1 and Channel 2.

**FIG. 2.**
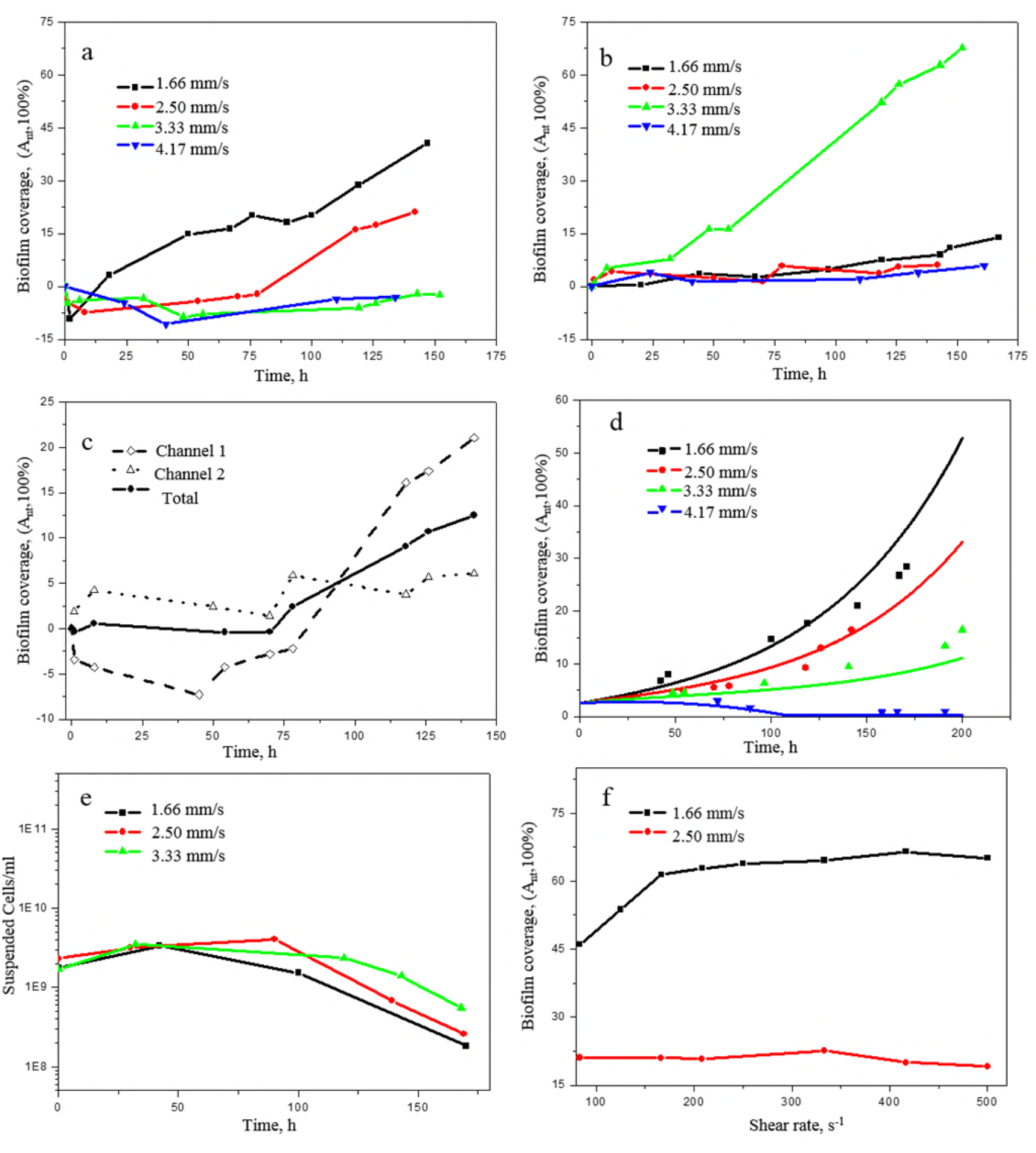
Optical images of biofilm growth in both microchannels at 1.0 N and various velocities. Images in the left column were taken after injecting nutrients for 1 h. The middle column shows images of biofilm growth for around 6 days. The right column lists images of biofilm detachment by increasing shear rate to 500.00 s^-1^. Nutrients flow from right to left in the upper channel. Scale bars indicate 100 μm.

### Biofilm accumulation in the flowing and no-flowing channels

Biofilm coverages as a function of time for different flow velocities in two microchannels are listed in Fig. 3. In the early of injection, the coverage of biofilms decreased as the flow shear stress snapped off some of weak initial attachments. After an active time when the left biofilms turned into irreversibly attached and new biofilms formed, biofilms coverage increased over time. As the velocity increased from 1.66 to 4.17 mm/s, biofilm coverage gradually decreased. Fig. 3 (b) plots biofilm coverage in the no flowing channel (Channel 2) as a function of time in each run. Biofilm coverages at all velocities increased over time, while the optimum velocity is 3.33 mm/s due to its exceptionally high accumulation rate.

**FIG. 3.**
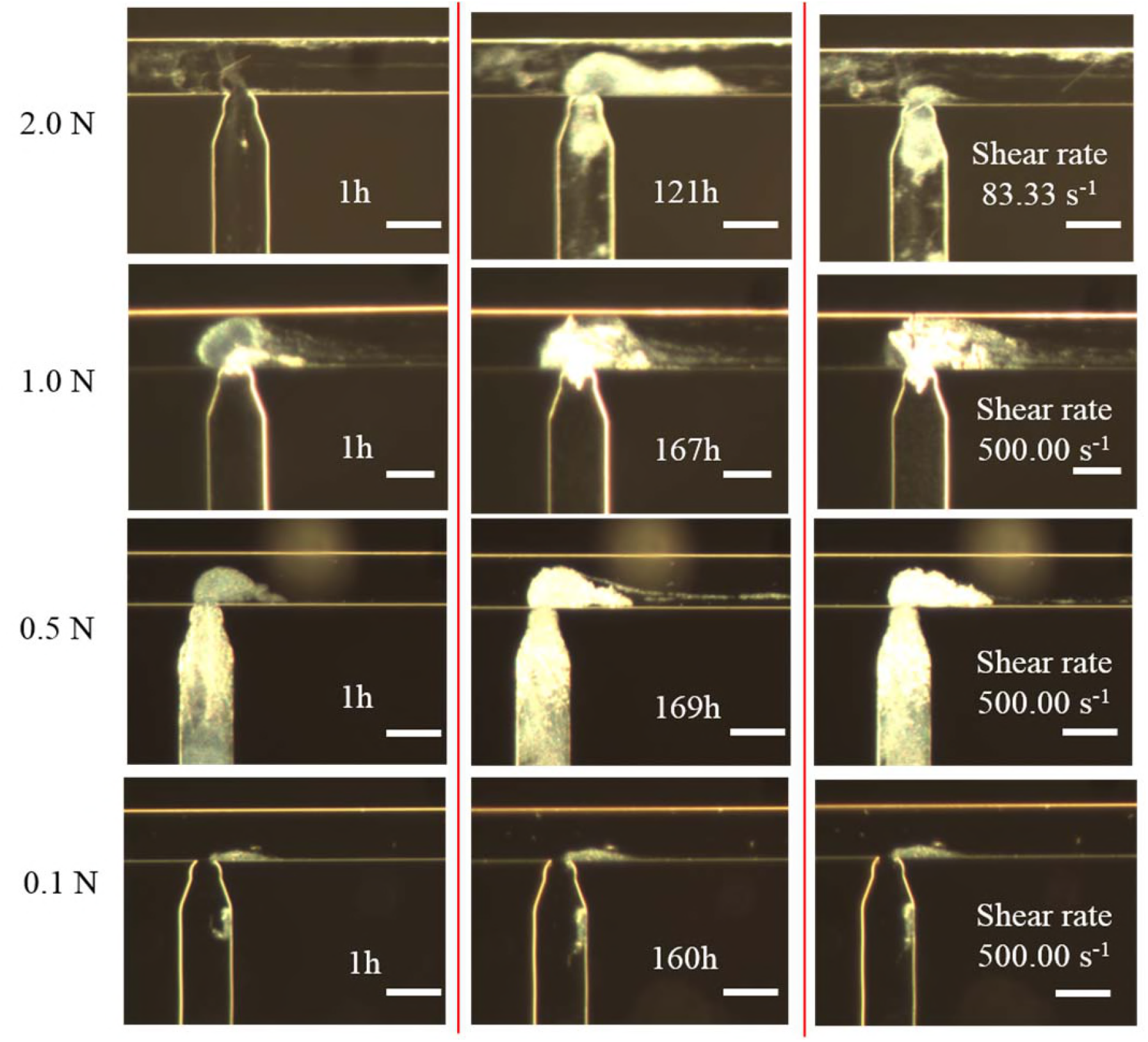
(a) Biofilm coverage over time in Channel 1 at various velocities; (b) Biofilm coverage over time in Channel 2 at different velocities; (c) Comparison of biofilm accumulation in both channels at 2.50 mm/s; (d) Experimental data and numerical simulations of biofilm coverage in both channels at various velocities; (e) Number of released cells as a function of biofilm culture time at various velocities; (f) Biofilm coverage in Channel 1 as response to the increasing shear rate after culturing biofilms at the velocities of 1.66 and 2.50 mm/s for 6 days. Comparing biofilms growth at 2.50 mm/s in Channel 1 and Channel 2 in Fig. 3 (c), biofilm coverage in Channel 2 increased with time initially but after 75 hours reached to a plateau value. This leveling off behavior was not observed in Channel 1. The stable coverage obtained in Channel 2 might be attributed to that cells in the biofilm cannot obtain sufficient essential sources of nutrients for new biofilm formation as biomass increased in the growing biofilm community. However, the continuous nutrients supply in Channel 1 delays this plateau. Fig. 3 (d) compares the experimental data with the mathematical model of biofilm coverages in both microchannels at various velocities. The numerical data is from D. L. Marbán’ work (D. L. Marbán, submitted for publication), and shows that our experiment data is well fit with the numerical simulation.

### Biofilm adhensive strenth test

In the biofilm culturing time, the number of released bacterial cells in the effluent at various velocities is shown in Fig. 3 (e). The cells number increased in the first two days after inoculation, which mainly contributes to that the reversible adhered bacteria after inoculation were driven out the microchannel by the nutrients flow shear stress. After 48 h, the biofilm-dispersal cells decreased, which corresponds to the increase of biofilm coverage over time in Fig. 3 (a), exhibiting that more bacteria involved into biofilm growth. Moreover, the increase of cell densities with flow velocity may indicate a higher detachment rate and a possible higher planktonic growth with an increase of shear stress.

After 6 days of biofilm culturing, biofilm coverages in Channel 1 as responds to the increasing shear rate from 83.33 s^-1^ and 125.00 s^-1^ up to 500.00 s^-1^ are shown in Fig. 3 (f). Biofilm accumulation at 1.66 mm/s increased when increasing nutrients shear rate to 166.67 s^-1^, suggesting that the increasing shear stress facilitates the diffusion of nutrients inside of biofilm and promotes its growth. Continuely increasing the shear rate, the growth trend slowed down; until up to 500 s^-1^, biofilm coverage slightly decreased, which dominates that the high shear rate brought about biofilm detachment. Simillar results are obtained at biofilm growth at 2.50 mm/s, which no large degree of detachment occurred as responds to low flow shear rates until up to 500 s^-1^.

### Effect of nutrient concentration on biofilm accumulation and adhesive strength

To assess the influence of nutrient conditions on biofilm accumulation and adhesive strength, biofilms were cultured at different nutrient concentrations. The baseline, 1.0 N, was 10 mM pyruvate in the growth medium and variations of two times (2.0 N), half (0.5 N) and one tenth (0.1 N) of the baseline concentration were applied. Injections were performed at a constant velocity of 1.66 mm/s from Channel 1 for approximately 7 days, and followed by a biofilm strength test by steadily increasing shear rate. The images are shown in Fig. 4.

### Biofilm morphologies

As shown in Fig. 4, biofilm in Channel 1 with the highest concentration 2.0 N has a long, thick but loose structure, which is highly sensitive to the variation of shear stress. After 122 h, the formed biofilm was dispersed from the deep of the matrix, leaving behind a few attached biofilm spots to regrow. At nutrients input 1.0

N and 0.5 N, biofilm became denser and compacted, and the influence of shear stress reduced. When decreasing the nutrient concentration to 0.1 N, there was no clear biofilm growth occurred in the nutrient continuous flowing channel.

The biofilm in Channel 2 at nutrient inputs of 2.0 N and 0.5 N had larger coverages than other concentrations, which the former confirms that high nutrient concentrations lead to a fast biofilm growth and the later might be related to the large initial attachments containing more biomasses for biofilm growth. It is noticed that there is barely new biofilm formation at both channels at 0.1 N, which shows that the lowest nutrient input significantly limited biofilm growth and formation.

As responding to the increasing shear rate, biofilm with low density and loose structure at 2.0 N, wars highly sensitive to the variation of shear stress, which detached from the substrates at the shear rate of 83.33 s^-1^. Biofilm growth at 0.5 N reacted as same as that at 1.0 N when the increasing shear rate acted on biofilms, and became elongated in the flowing direction.

**FIG. 4.**
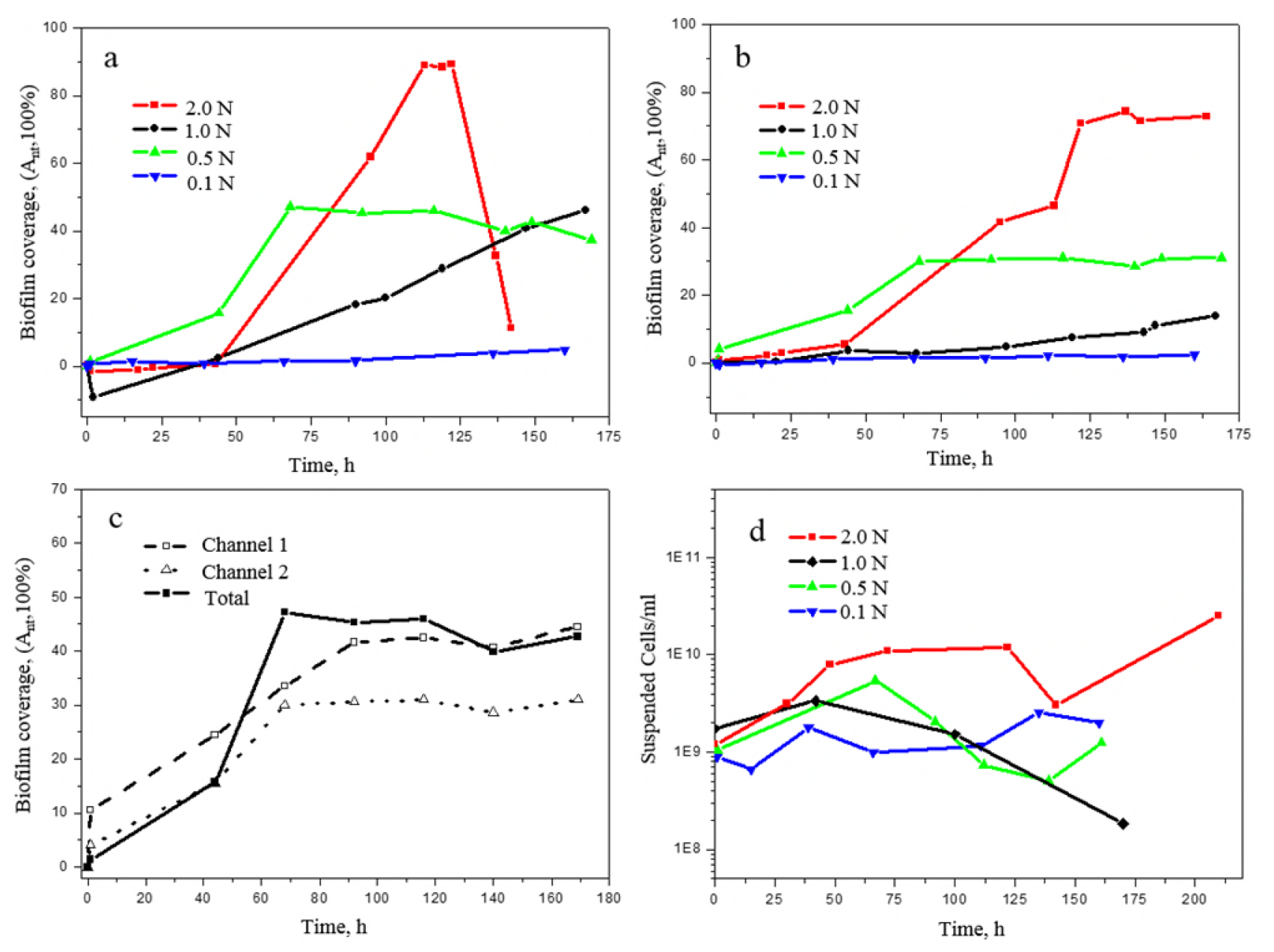
Optical images of biofilm growth over time at various nutrient concentrations, 2.0 N, 1.0 N, 0.5 N, and 0.1 N, respectively. Images in the left column were taken after injecting nutrients for 1 h. The middle column shows images of biofilm growth for around 7 days. The right column lists images of biofilm detachment by increasing shear rate to 500.00 s^-1^. Nutrients flow from right to left in the upper channel. Scale bars indicate 100 μm.

### Biofilm accumulation in the flowing and no-flowing channels

Biofilm coverages as a function of time for different nutrient concentrations in two microchannels are shown Fig. 5. As shown in Fig. 5 (a), in Channel 1, biofilm growth at a high nutrient concentration of 2.0 N has a much faster accumulation rate in the first 5 days, but rapidly decreased when most parts of biofilm matrix were detached from the substrate. At the medium nutrient feeding zones, biofilm accumulation at 0.5 N is higher than that of 1.0 N in the first 3 days, and reached a plateau value after that, which was not observed for 1.0 N. When decreasing the nutrient concentration to 0.1 N, there was no clear biofilm formation in both channels. Therefore, the lowest nutrient concentration (0.1 N) could not provide environment for biofilm growth. In this study, the limiting nutrient concentration for biofilm growth appears to be between 0.1 and 0.5 times N.

**FIG. 5.**
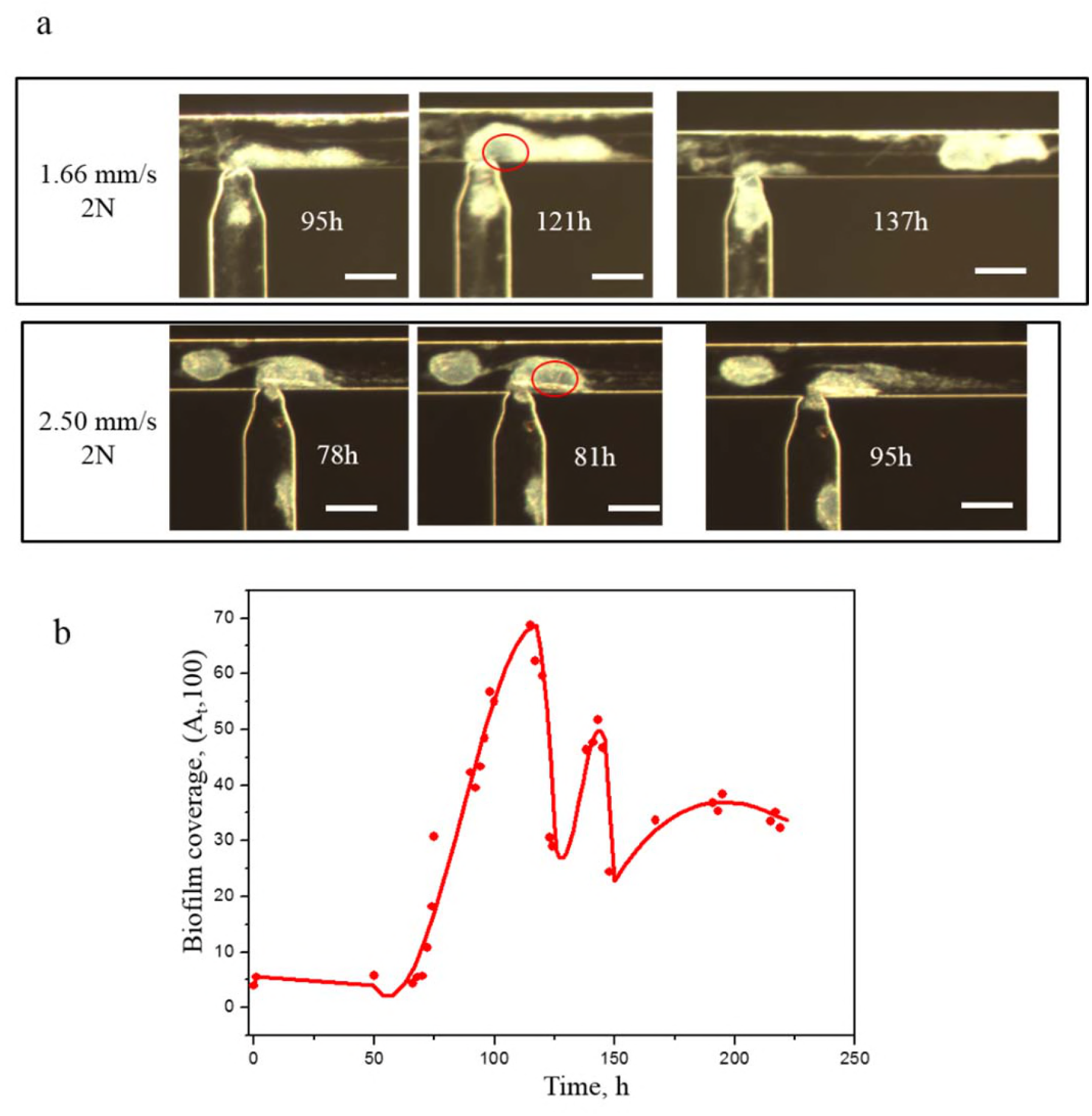
(a) Biofilm coverage over time in Channel 1 at different nutrient concentrations; (b) Biofilm coverage over time in Channel 2 at different nutrient concentrations; (c) Comparison of biofilm coverage in both channels at 0.5 N and 1.66 mm/s; (d) Cell number of effluents at various nutrient concentrations at the flow velocity of 1.66 mm/s. As shown in Fig. 5 (b), biofilm accumulation in Channel 2 is influenced by nutrient concentrations. Biofilm formation at 2.0 N has larger coverage than other cases, indicating that high nutrients loading in Channel 1 leads to an increase in biofilm growth in Channel 2. In addition, biofilm growth in a no flow channel reached to a stable plateau at the later stage of its development. The time to reach the stable plateau at 2.0 N was later than 0.5 N, suggesting that high nutrient concentration leads to a decrease in the time taken to reach the stable plateau in a no flow system. Fig. 5 (c) shows that biofilm coverage obtained stable plateaus at 0.5 N in both channels. The time to reach the plateau in Channel 1 was later than that in Channel 2, indicating that flow shear rate can facilitate mass transfer and lead an increase in the time taken to reach the stable state.

### Biofilm adhesive strength test

**Fig. 5 (d)** presents the result of cell number in the effluent at different nutrient concentrations. The cell number at 2.0 N is higher than other nutrient concentrations. The released cell numbers are relatively in the same level at 0.5 N and 1.0 N in the beginning. However, when biofilm stopped growing at 0.5 N, the detached cells increased over time, suggesting that the mature biofilm would disperse more planktonic cells into the bulk liquid (26). At the limited nutrient supply (0.1 N), the released cell number in the effluent was stable and no biofilm accumulated in the channel, indicating that bacteria at limited nutrient loading prefer to live in the planktonic style instead of biofilm style (36, 37).

It is noticed that biofilm growth at 2.0 N had a weak adhesive strength with substrates, because cells deep in the biofilm were dispersed from the interior of the biofilm matrix causing large degree of detachment. We observed this dispersion occurring at nutrient concentration of 2.0 N and flow velocities of 1.66 and 2.50 mm/s (Fig. 6 (a)). Firstly, a central region in the biofilm matrix (shows in red circles of images in Fig. 6 (a)), become visible and light, which has demonstrated the pre-dispersion behavior (22). Eventually, microcolonies within the regions migrated into the bulk liquid, leading to huge biofilm detachments. Biofilms were observed to undergo growth and dispersion simultaneously at high nutrient concentrations (Fig. 6 (b)). The coverage area increased steadily after an active time, but decreased when biofilms were detached from the substrate and increased again while the left biofilm spots regrew. As biofilm growth at high rate at 2.0 N, cells trapped deeper in the biofilm matrix may have difficulties obtaining essential sources of energy or nutrients. In addition, waste products and toxins can accumulate fast in the biofilm community to reach toxic levels, threatening cells survival. Thus, microorganisms within the biofilm release from the matrix to resettle at a new location.

**FIG. 6.**
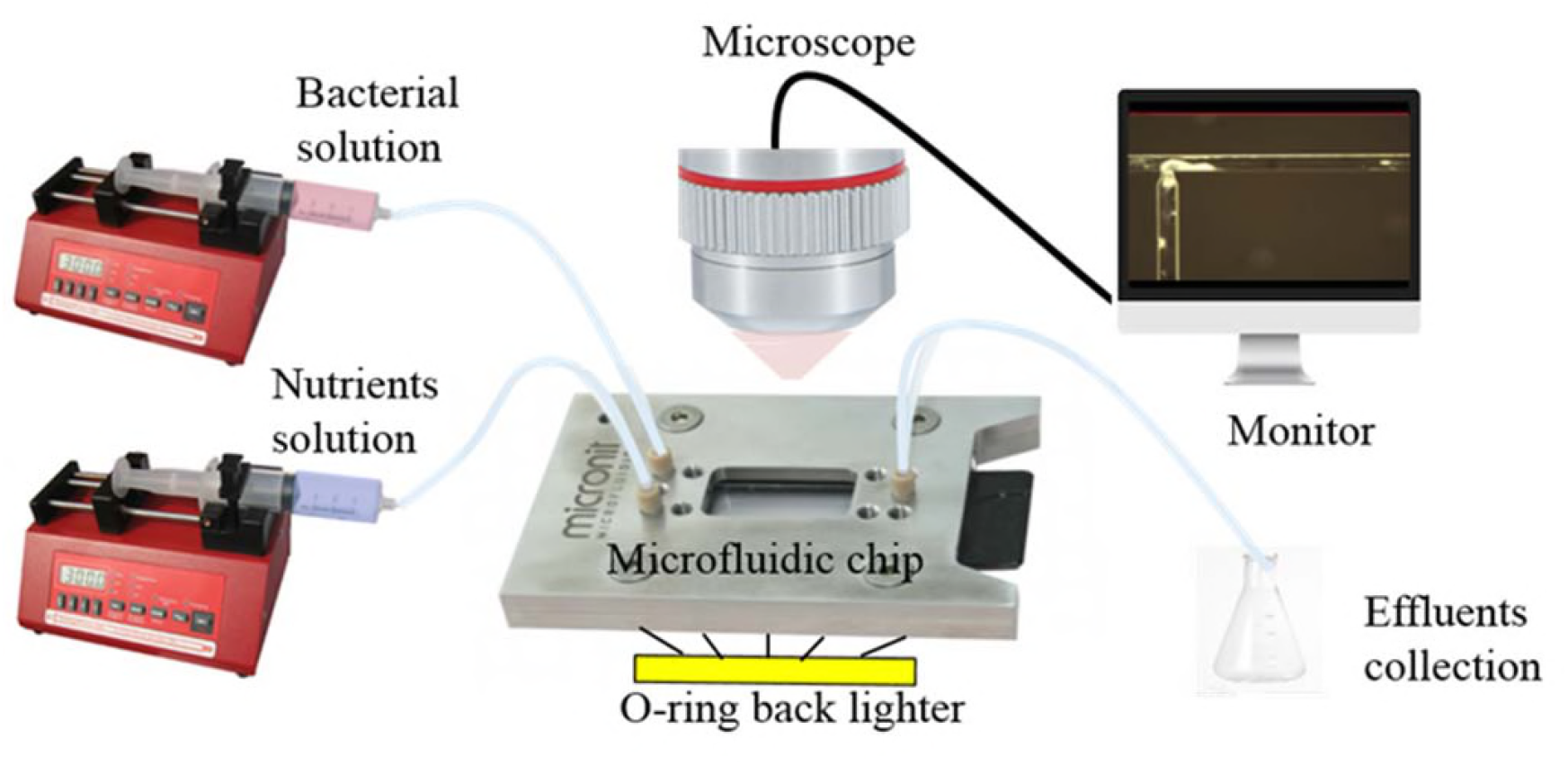
(a) Images of biofilm growth following dispersion events at high nutrient concentration of 2.0 N and flow velocities at 1.66 and 2.50 mm/s. (b) Biofilm accumulation at 1.66 mm/s and nutrient concentration of 2.0 N.

## DISCUSSION

### Biofilm morphologies

The observations on biofilm morphologies at each run demonstrate that flow velocity and nutrients concentration have direct effects on biofilm morphology. Biofilms formed in Channel 1 reveal the influence of flow shear stress drag.

The shapes of biofilm clusters became compacted and progressively elongated along the flow direction with the increase of flow velocity (Fig. 2). While biofilms formed at the high nutrient concentration have long, thick but loose structures, and became denser and compacted with the decrease of nutrient loading (Fig. 4). Similar results have been reported in previous work (21).

Biofilm growth in Channel 2 is highly dependent on the diffusion of nutrients in Channel

1.As the former bacteria injection path, most parts of Channel 2 were full of biomasses without fluid shear forces. Only the void in the nozzle connecting with Channel 1 could act as the transport channel supplying nutrients for biofilm growth. Biofilms at the high shear rate of 166.67 s^-1^ and 2.0 N led to larger clusters compared with others, indicating that shear rate and nutrient concentration in Channel 1 determined the flux of nutrients transport to Channel 2.

It is noticed that there was no biofilm growth in either channel at the highest flow velocity of 4.17 mm/s and lowest nutrients concentrations of 0.1 N, suggesting that the high shear forces and limited nutrients loading may prevent biofilm formation, which is in agreement with industrial applications where the formation of biofilm is prevented by high velocity flooding (36).

### Biofilm accumulation in the microchannel

In this study, we set the initial biofilm coverage after inoculation to zero, and plot biofilm net coverage A_nt_, by subtracting initial attachment to analysis biofilm accumulation. As shown in Fig. 3 (a) and Fig. 5 (a), the coverages of biofilm are under zero in Channel 1 in the early stage of injection, which demonstrates that the shear stress caused by nutrients flowing leads to snap-off of weak initial attachments. When the remained biofilms became irreversibly attached, they behaved as nuclei for new bacteria/biofilm growth, resulting in the increase of biofilm coverage. Biofilm accumulation in the flowing microchannel (Channel 1) is highly related with flow velocities through two important factors, mass transfer and shear stress (19, 27). As shown in Table 1, the Reynolds numbers in Channel 1 were very low (from 0.17 to 0.42), while the mass transfer Peclet number were extremely high (from 97.64 to 245.30), which suggests that mass transfer in the microchannel was dominated by convective actions and has negligible diffusion (38). Thereby, the diffusion of nutrients from bulk to biofilms rarely increased when increasing the flow velocity, while the shear stress caused by water flow increased linearly. The accumulation of biofilm, which is equal to its growth rate minus detachment rate, decreased with increasing flow velocities when the shear stress induced detachment rate exceed growth rate. Thereby, the optimum flow velocity for biofilms growth in the flow microchannel is the lowest velocity of 1.66 mm/s in this work.

The effect of nutrient concentration on biofilm accumulation in Channel 1 is a non-linear relationship. The observations at 0.5 N and 1.0 N implies that, in a range of concentration of nutrient, the biofilm growth rate is independent of the nutrient concentration in the beginning of biofilm growth (29); as biofilm growing in size, biomass demand is rising steadily, thereby the nutrient concentration determinates the growth rate in the later stage of biofilm development.

Biofilm accumulation in Channel 2 increased with shear rate and nutrient concentration in Channel 1 monotonically. Due to in absence of shear stress, biofilm growth in Channel 2 depends on the nutrient diffusive flux from Channel 1, which increases with the flow velocity and nutrient concentration. Therefore, for a confined no flowing system, biofilm accumulation rate is highly related to the nutrients availability, while the flow shear rate facilitates mass transfer, leading to an increase in biofilm accumulation. These observations are in correspondence with previous works (20, 21, 37).

The results indicates that for porous systems, like oil reservoirs, biofilm could develop not only in the main water flow paths, but also in dead ends and less flooded areas. Therefore, optimized nutrient flow velocity and nutrient concentration ensures sufficient nutrients supplying rate with moderate shear stress in the microchannel, resulting in biofilm accumulation in both flowing and non-flow regions.

### Biofilm adhesive strength with the glass surface

Since only the nutrients solution was injected through Channel 1 after bacterial inoculation, the suspended cells in the effluent can be interpreted as the detachment of biofilms which dispersed their planktonic cells in the bulk growth medium. During exposure to stress, including shear stress and nutrient starvation, cells dispersed from biofilms go into the planktonic growth phase (39, 40). In this study, we observed that the biofilm-dispersal cells increased with flow velocity due to the shear stress induced detachment; nutrient starvation was also a trigger for biofilm dispersal. In addition to poor nutrient loading, biofilms culturing at the high nutrient concentration (2.0 N in this study) could also result in biofilm dispersal in the deep of biofilm matrix, which is mainly because that cells trapped deeper in the biofilm matrix may have difficulties obtaining essential sources of energy or nutrients. In a flowing system, biofilm dispersal is beneficial to spawn novel biofilm development cycles at new locations. Therefore, biofilm dispersal can potentially be used to control bioplugging in the further places of porous media.

In contrast to the planktonic mode, biofilm in a self-generated matrix can behave as viscous liquids to resist the flow shear stress and prevent from detachment from the attached solid surface. The results from biofilm adhesive strength test have demonstrated that biofilms growing at medium nutrient concentrations (0.5 N and 1.0 N) could resist the flow-induced shear stress. Compared to the large detachment at the initial stage, it suggests that the adhesive strength between biofilms and adhesive surface became stronger under shear (22, 41, 42). However, biofilm growth at high nutrient concentration (2.0 N) forms a loose structure with a high accumulation rate but a weak adhesive strength with substrates, which is easily detached by fluid shear.

In conclusion, this work demonstrates that flow velocity and nutrient concentrations can have significant impacts on biofilm development in both flowing and stagnant microchannels. Negligible biofilm formation at the relatively high flow velocity of 4.17 mm/s and low nutrient concentration of 0.1 N suggests that there is a ‘no/low growth region’, where high shear forces lead to biofilm detachment and nutrient concentration is below the minimum required for biofilm formation. This is supported by the earlier work (21). At the conditions investigated in this work, a strong plugging effect in the flowing microchannel was obtained at the relatively low flow velocity of 1.66 mm/s and the medium nutrient concentration of 1.0 N (10 mM substrate), which has a relative fast biofilms accumulation rate and a strong adhesion force to resist increase in the flow-induced shear. This research gives new insight to the relative influences of flow velocity and nutrient concentration on biofilms development at pore scale. This may aid evaluations of bioplugging in porous systems such as for oil and ground water reservoirs. As potential permeability reducers in oil reservoirs, biofilm accumulation in porous media needs to be controlled by flow velocity and nutrient availability. Optimized nutrient flow velocity and concentration ensures sufficient nutrients supplying rate with moderate shear stress in the microchannel, resulting in biofilm accumulation in both flowing and non-flow regions. However, too high stress may prevent biofilm formation and removal of adhered biofilms in the porous media. High nutrient concentration is beneficial for biofilm growth, but leads to a weak biofilm adhesive strength, which is easily detached by flow shear from the pores.

## MATERIALS AND METHODS

### Bacteria and fluids

The bacteria used in the study was:*Thalassospira strain A216101*, a facultative anaerobic, nitrate-reducing bacteria (NRB), capable of growing under both aerobic and anaerobic conditions. It is able to grow on fatty acids and other organics acids as sole carbon and energy source. Bacteria were enriched in a marine mineral medium, which contained the following components (L^-1^): 0.02 g Na_2_SO_4_, 1.00 g KH2PO4, 0.10 g NH4Cl, 20.00 g NaCl, 3.00 g MgCl2·6¾O, 0.50 g KCl, 0.15 g CaCh·2¾O, 0.70 g NaNO3, and 0.50 ml 0.20% resazurin (43). The medium is hereafter referred to as growth medium. After autoclaving in a dispenser, 1 L of growth medium was added 5 ml vitamin solution and 20 ml 1 M NaHCO3 to adjust the pH to 6.80-7.20. Finally, pyruvate was added as the carbon source from a sterile stock solution to achieve final nutrient concentrations of 20 mM (2.0 N), 10 mM (1.0 N), 5 mM (0.5 N), and 1 mM (0.1 N), respectively. The final nutrient medium was stored at 4°C.

### Experimental setup

The experimental apparatus is illustrated in Fig. 1. A T-junction microfluidic device (Micronit, Netherland) consists of a single straight channel and a side channel with the sizes of 100 μm width and 20 μm depth and the nuzzle size at the crosssection as narrow as 10 μm (Fig. S1). Two syringe pumps (NE-1000 Series of Syringe Pumps, accuracy ±1%) were used to load the bacterial inoculation solution and nutrients solution separately into the microchannels. The light source is a cold halogen lamp with 24v, 150w placed under the microchip for better illumination. The micromodel was then placed under a microscope with a digital camera (VisiCam 5.0, VWR) to acquire image sequences. Measurements and experiments were conducted at room temperature and pressure.

**FIG 1.**
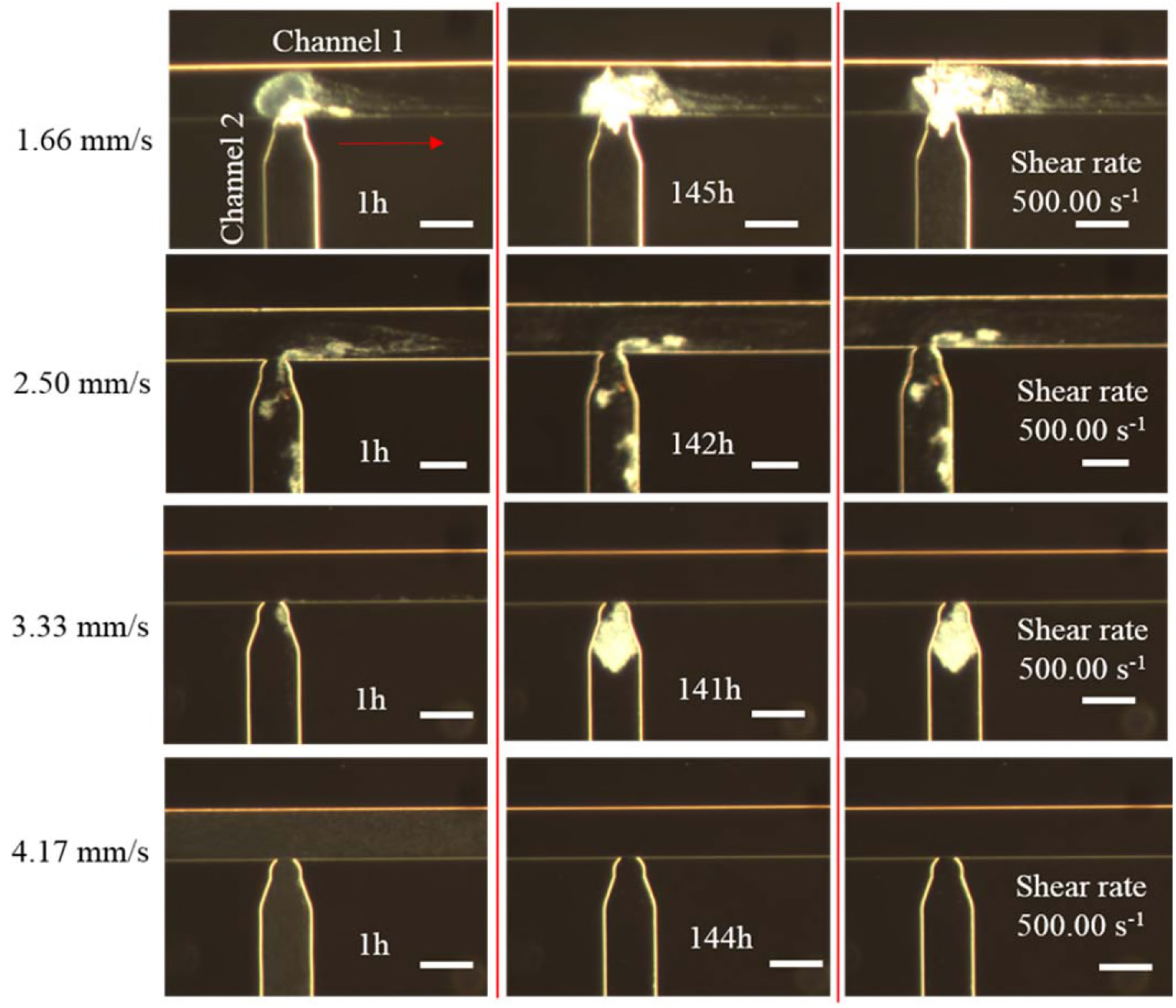
Schematic illustration of the experimental setup.

### Inoculation process

Before inoculation, the microchannel was cleaned using ethyl alcohol, deionized water, H_2_O_2_ solution (10 wt%) and deionized water to guarantee the same surface condition for each experiment. The bacterial inocula were pre-cultured in the growth medium containing 10 mM (1.0 N) nutrients at 30 °C for 24 h. Inoculation was achieved by injecting the pre-culture bacterial solution from the bacterial inlet port (Fig. S1 (a)) into the side channel at the rate of 1.0 μl/min for 24 h, followed by a 24 h shut-in period. In case of biofilm clogging the straight channel for nutrients injection, we separated the bacterial injection channel (Channel 2) and nutrients flowing channel (Channel 1) and closed the nutrient inlet during inoculation to force bacterial solution to only flow towards the outlet direction. Then only nutrients with various pyruvate concentrations (from 0.1 N to 2.0 N) were injected from the nutrients flow channel (Channel 1) at constant flowrates from 0.2 to 0.5 μl/min for approximately 6-7 days, while Channel 2 was closed, which led to a greater growing of bacteria on the substrates of the intersection of straight channel and side channel. Before the next experiment, microchannels were rinsed with ethyl alcohol, water, H_2_O_2_ solution and water separately, finally, filled with the marine medium without nutrients until the onset of the next experiment.

### Image process

Image sequences on biofilm growth were acquired with a Leica microscope fitted with a digital camera for scoring with time. The main area of interest in this study is the intersection of straight channel and side channel, thereby two areas of interest (AOIs) with 0.5mm*0.1mm are extracted from the origin image for further image analysis (Figure S1 (b)). The image processing was performed using MATLAB®’s Image Processing Toolbox. Biofilm accumulation, here presented by biofilm coverage (A_n_t) in areas of interest, was periodically measured in a flowing channel (Channel 1) and no-flowing channel (Channel 2). Further details on image process can be found in Support Information S1.

### Effluent PCR analysis

Fluid samples were collected daily at the outlet through a quantitative real-time PCR (qPCR) on whole-cells to determine the total number of bacteria. A 20 μl qPCR reaction mix containing 10 μl SYBR^®^ Green PCR kit, 0.06 μl primers (100uM), 8.88 μl nuclease free water and 1 μl template was made. The reaction was run by the following cycling conditions: denaturation of DNA at 95°C for 15 minutes, 36 cycles with denaturation for 30 seconds at 94°C, annealing for 30 seconds at 55°C, extension for 1 minute at 72°C followed by a plate read. At the end, a melting curve from 55°C to 95°C was conducted. The reactions were carried out in a CFX connect™ real time PCR detection system (BioRad).

## ACKNOWLEDGMENTS

We wish to thank Edin Alagic, Rikke H. Ulvøen and Tove L. Eide for technical assistance. This work was supported by the Research Council of Norway and industry partner GOE-IP through the projects IMMENS no. 255426.

